# Structural Motifs, Disorder, and the Efficacy of Viral Vaccines

**DOI:** 10.1101/2020.06.08.139907

**Authors:** Robert A. Makin, Steven M. Durbin

**Author notes:** These authors contributed equally to this work.

## Abstract

We demonstrate that it is possible to draw direct numerical correlations between virus particles and effective virus-like particle (VLP) derived vaccines through extraction of a Bragg-Williams order parameter from electron microscopy. The method has its roots in studies of disorder in metal alloys, and is adapted to describe the type and occurrence of structural motifs within the arrangement of viral coat proteins, captured by the value of the order parameter as a measure of disorder. A conventional approach to viral vaccine design consists of replicating select proteins to create a VLP designed to trigger an immune response while remaining non-infectious. Understanding variations between viruses and vaccine strains therefore tends to focus on differences between proteins, which can be characterized through genetic analysis. While such an approach provides vital information about the functioning and interactions of the proteins, it does not yet yield an early-stage pathway towards predicting the efficacy of a vaccine, and so large-scale clinical trials are required to obtain critical information. With the urgency associated with pandemics, including Coronavirus Disease-2019 (COVID-19) originating from the SARS-CoV-2 virus, there is a need for earlier indications of whether a vaccine has the necessary characteristics. Application of the methodology to Dengue and influenza virus particles indicates that temperature and pH during incubation could potentially be exploited to fine-tune the order parameter of VLP-based vaccines to match the corresponding virus. Additionally, utilization of an Ising model plot reveals a clear relationship between case fatality rate and order parameter for distinct virus families.

## Introduction

There is considerable interest in reducing the time necessary to develop vaccines, especially in light of the Coronavirus Disease-2019 (COVID-19) pandemic stemming from the SARS-CoV-2 virus [1]. Here, we demonstrate that what might be best described as a hidden variable – when properly recognized – identifies a necessary characteristic for any effective vaccine derived from virus-like particles (VLPs). The underlying concept relates to what are known as structural motifs, which are commonly used to understand proteins and their behavioral variations [2]. The notion of a structural motif is, in fact, quite general, and can be employed to describe spatial arrangements of proteins – such as what occurs on the surface of a virus particle – extending the concept beyond variations within the protein structure itself. The advantages of such an approach can be significant, as it is the disorder and relative composition of viral coat proteins which characterizes such surface configurations that must be preserved in vaccines. Further, the quantifiable degree of disorder can be used in the context of an Ising model to predict the case fatality rate of a new virus, and even dictate the environmental process parameters to monitor for incubation, such as pH and temperature, for potential vaccine components.

## Discussion and results

A common structural feature of viruses is the outer-most layer of proteins, which are responsible for activities such as binding to host cells and injecting viral genetic material into those cells. The outer proteins (referred to as viral coat proteins) also play an important role in the immune system response, since it is to these proteins that antibodies bind in order to signal the immune system to attack the virus. It is possible to view the configuration of these outer-most proteins from a structural motif perspective, as they are typically six-fold coordinated (although other coordination numbers are possible), with a central protein surrounded by six other proteins to form the basis of a hexagonal-close packed lattice structure (Fig. 1). There are numerous possible permutations of six-fold motifs. In the case of a virus with two major viral coat proteins (which, to avoid loss of generality, we will refer to as *α* and *β*) as depicted in Fig. 1A, there are 20 distinct six-fold motifs. The number of each motif occurring in a particular specimen is directly determined by the viral coat protein composition and by the degree of disorder characterizing the virus surface, the latter of which can be quantified through a Bragg-Williams type order parameter *S* [6,7]. A fully ordered surface, corresponding to a unity value of *S*, consists of only a single structural (base) motif, illustrated in Fig. 1 a, b and c for the cases of six-fold coordinated viruses with two and three major viral coat proteins, respectively. In contrast, a completely disordered protein configuration corresponds to a value of *S* = 0, and can have a large set of structural motifs representing variations of the base motif.

**Fig 1.**
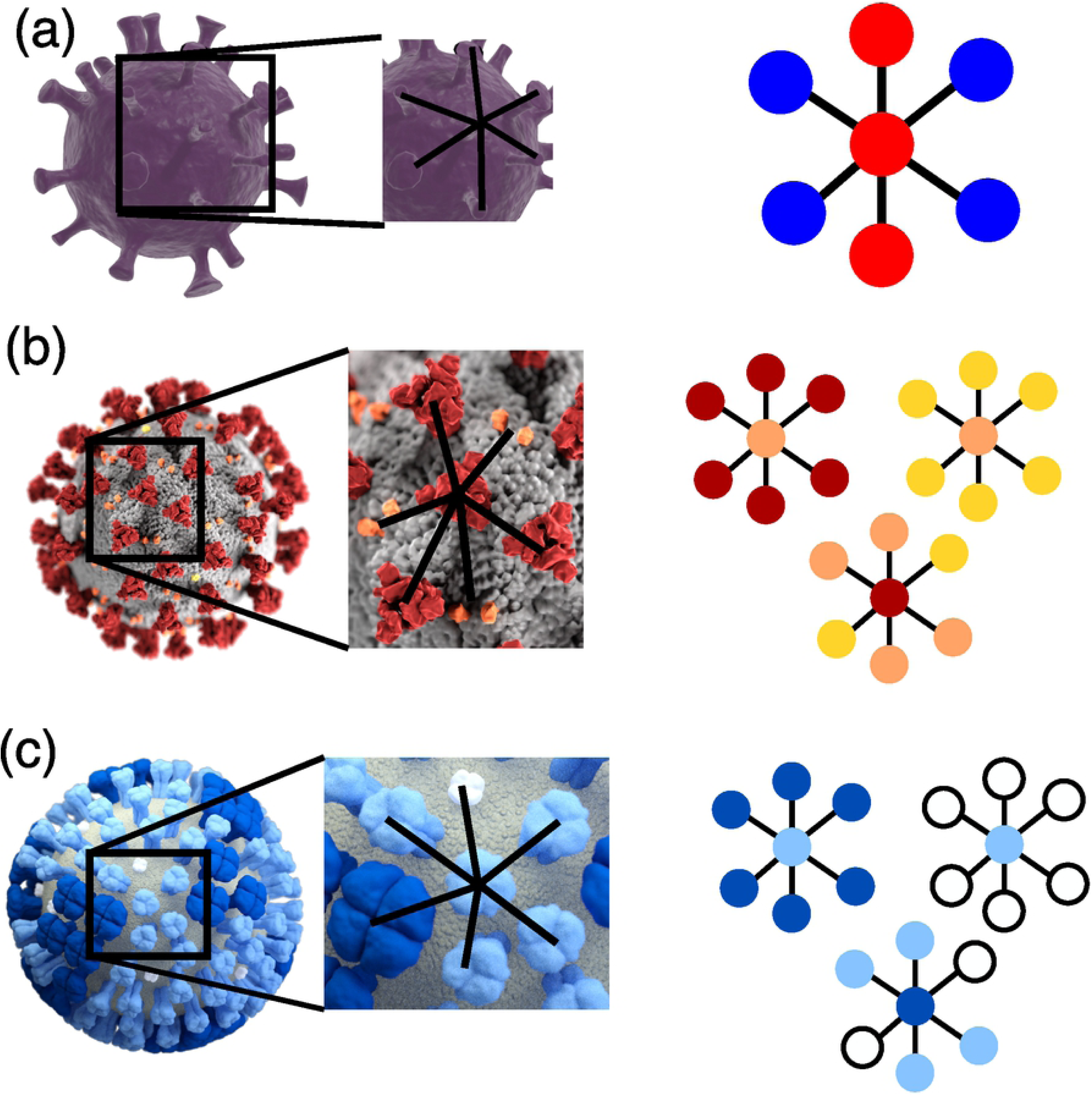
Illustration of surface protein motifs. (a) Henipavirus particle with six-fold motif highlighted, model created using ultrastructure description from [3]. Base motif is shown schematically with fusion protein (blue) and glycoprotein (red). (b) Beta-coronavirus particle. Adapted from [4]. The three base motifs are shown schematically with spike protein (red), membrane protein (orange) and envelope protein (yellow). (c) Influenza virus particle. Adapted from [5]. The three base motifs are shown schematically with the M2 ion channel in white, hemaggluttinin in light blue and neuraminidase in dark blue.

One of the two major factors that determine the occurrence probability of the different motifs is the ordering of the viral coat proteins on the outer surface of the virion. To proceed with developing a quantifiable measure of the degree of ordering, we first define the perfectly ordered (*S* = 1) case. This is accomplished by selecting the base motif – the structural motif which contains equal numbers of both viral coat proteins, identified in Fig. 1B (note that for this case the outer viral coat proteins of the motif only contribute ½ to the viral coat protein count since they are shared by neighboring motifs). Using the base motif as a reference, the degree of ordering can be quantified through the Bragg-Williams long range order parameter *S*, where *S* = *r*_*α*_ + *r*_*β*_ – 1. Here, *r*_*α*_ is the fraction of *α* viral coat proteins on *α* -sites, and *r*_*β*_ is the percentage of *β* viral coat proteins on *β*-sites, respectively (from the perspective of the reference motif); it is straightforward to extend the approach for situations involving more than two distinct viral coat proteins. In material systems such as binary metal alloys or semiconductors, the order parameter historically has been measured through techniques such as x-ray diffraction [7]. Recently, we have extended the determination of *S* to include techniques such as Raman spectroscopy [8] and electron microscopy [9, 10], the latter of which we have used to calculate the order parameter of virions from transmission electron microscopy (TEM) images.

The other major factor that determines the occurrence probability of different motifs is the relative composition of the viral coat proteins. For the case of two such proteins, we define composition as 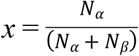, where *N*_*α*_ and *N*_*β*_ represent the total number of each protein, respectively, and can be measured through techniques such as sodium dodecyl sulfate–polyacrylamide gel electrophoresis (SDS-PAGE) [11]. The percentage of different motifs that will occur on the surface of the virion can be calculated based on the composition *x* and the order parameter *S*. Figure 2 shows the motif percentages for a fully disordered surface over the full range of possible compositions of a two-viral coat protein, six-fold coordinated motif. As illustrated in Fig. 2 a, the base motif and its related variations are much more likely to form at balanced compositions, i.e., near *x* = 0.5. At the extreme values of *x*, the motifs dominated by a single viral coat protein (β_7_ and α_7_ motifs) become much more likely to form, and specific types of single viral coat protein dominated motifs (such as the α_2_β_5_ or α_6_β_1_ motifs) become more likely with shifts towards the extreme ranges of the composition, as can be seen in Fig. 2 b. We note that depending on the composition, there may be physically inaccessible values of *S*; the maximum *S* value for *x* ≤ 0.5 is 2*x*, and 2(1-*x*) for *x* ≥ 0.5.

**Fig 2.**
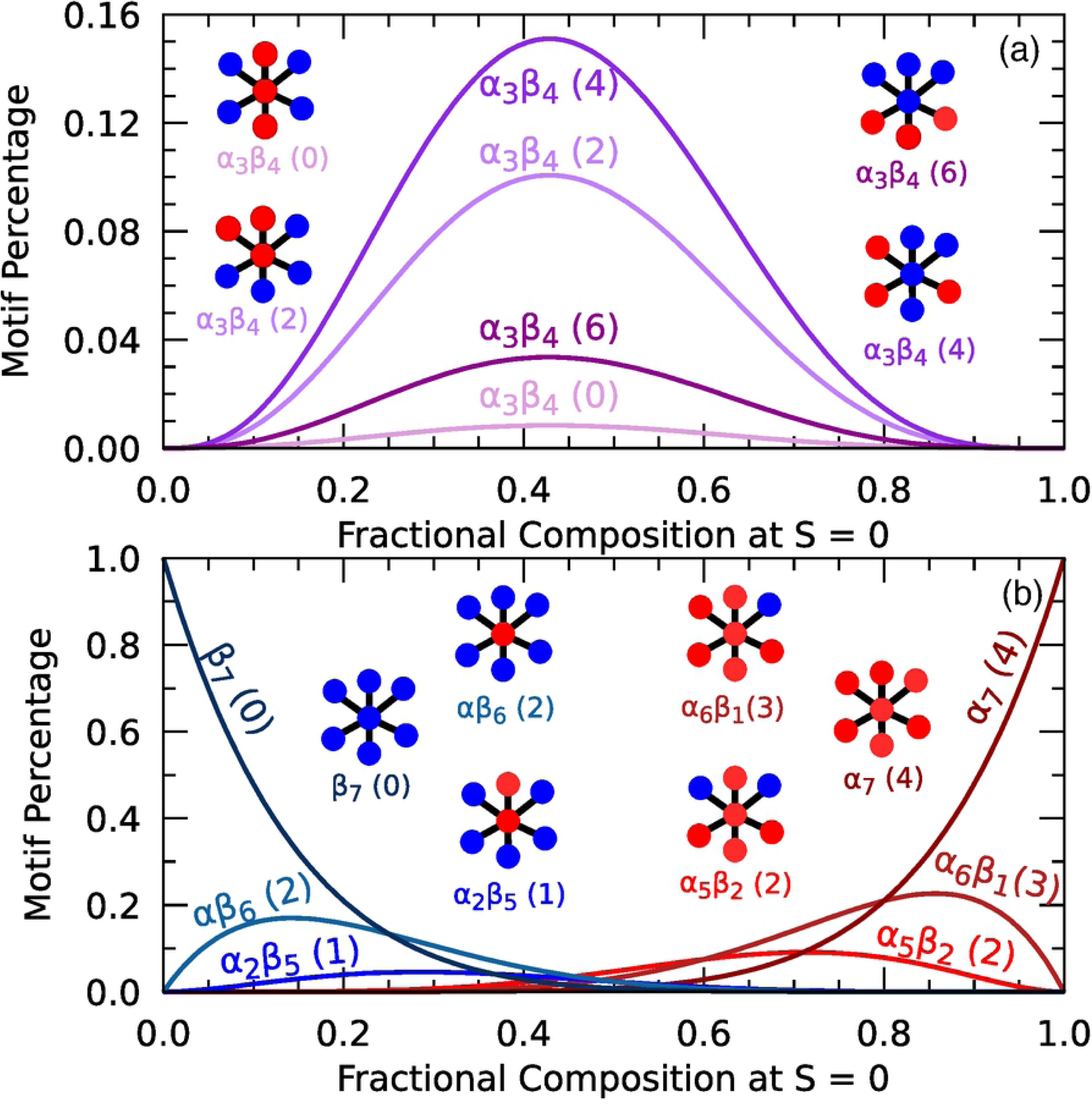
Percentage occurrence of different structural motifs at *S* = 0 as a function of composition for two viral coat proteins. (a) α_3_β_4_ base motif along with the complete set of variations; (b) Representative selection of single-viral coat protein dominated motifs. The number in parenthesis indicate the number of proteins on antisite positions for that motif.

Since viral coat proteins play a defining role in the interactions between viruses, host cells, and antibodies, and the viral coat protein structural motifs play a significant role in these interactions, we propose that when developing vaccines starting with VLPs, it is important to match the motif composition of the intended virus. Counting the motifs of a given virus or VLP is impractical, however, and so taking advantage of the correlation between the numerical occurrence of specific motifs and the order parameter for known viral coat protein composition provides a viable alternative. Figure 3 compares order parameter values extracted from TEM images for five distinct viruses to the corresponding experimentally determined order parameters for VLPs used to create vaccines. The VLPs for the two influenza viruses and for the Zika virus have order parameters very close to their corresponding viruses, and showed promising results for developing immune responses. The VLP shown for HPV16 corresponds to Gardisil®9, one of two commercially available vaccines. Conversely, for the example HIV vaccine in Fig. 3, the VLP has an order parameter much higher than that of the virus. A possible explanation for the difficulty in developing an effective HIV vaccine [21,22] could therefore be that current vaccines and even potential vaccines, such as yeast based VLPs [23] (the *S*^2^ of one such yeast-based VLP is shown in Fig. 3), do not properly match the order parameter and composition [24] of the viral coat protein surface of the HIV virus. This suggests that for a vaccine to be effective, the order parameter of any VLP used in its creation should closely match that of the corresponding virus.

**Fig 3.**
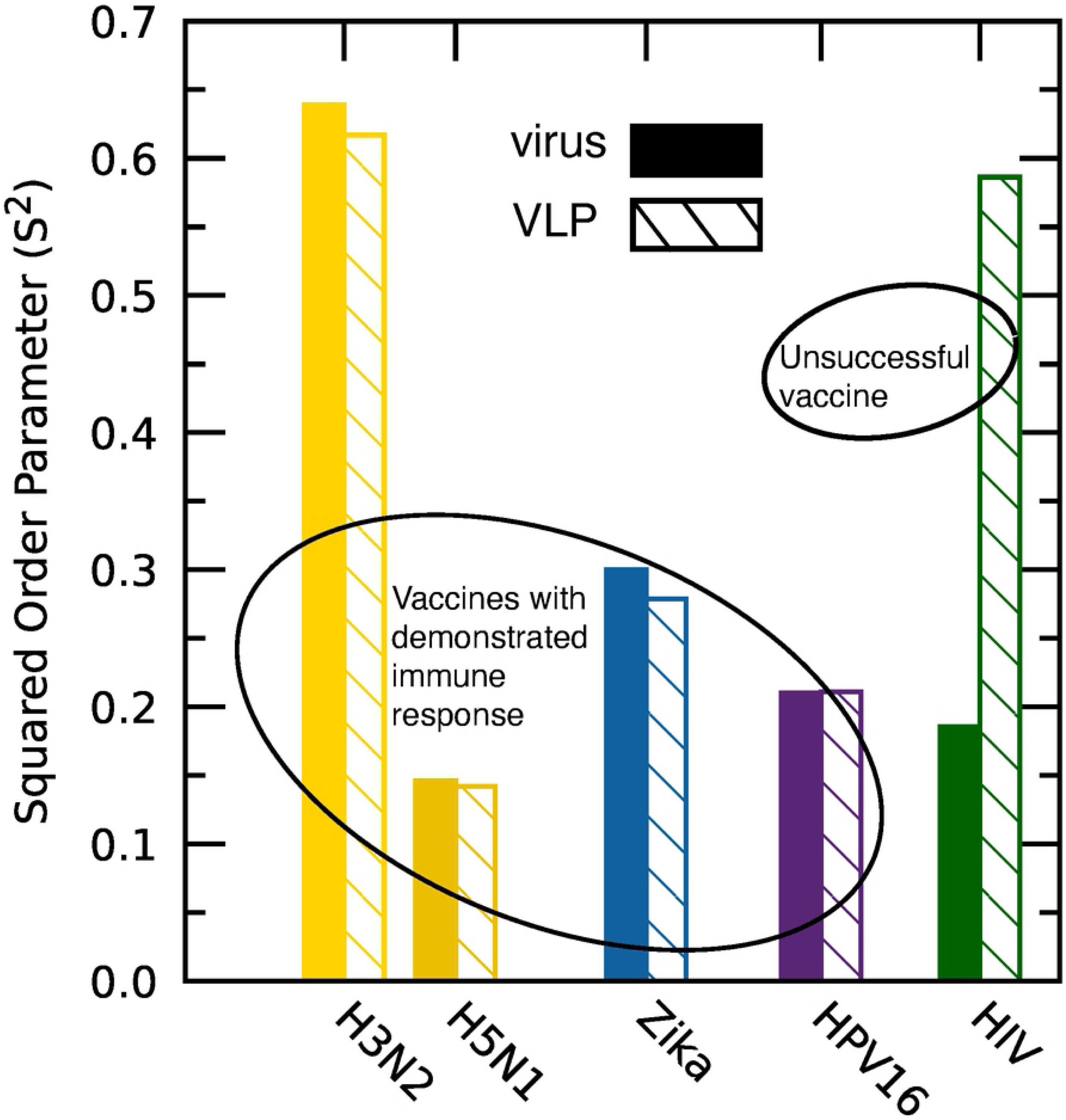
Comparison of order parameter (plotted as *S*^2^) for selected viruses and their corresponding VLP. Data extracted from TEM images obtained from H3N2 [12] and H5N1 VLPs [12]; H3N2 Virus [13]; H5N1 Virus [14]; Zika virus [15], VLP [16]; HPV16 virus [17], VLP [18]; HIV virus [19], VLP [20].

When developing VLPs (or any vaccine), the composition of the viral coat proteins can be controlled by gene manipulation. However, the degree of disorder which characterizes the viral coat protein surface of a virus is influenced by conditions under which the virion matures; for example, the temperature or pH of the growth environment. This is observed in the plot of Fig. 4, which shows the variation of the order parameter in Dengue and influenza viruses with varying incubation temperatures in a neutral pH environment. The readily apparent linear variation of the square of the order parameter with temperature is indicative of a Landau second-order phase transition present in ordered-disordered systems [25]. Additionally, the inset shows the influence of pH on the ordering of the Dengue virus, which follows a similar linear trend as the temperature.

**Fig. 4.**
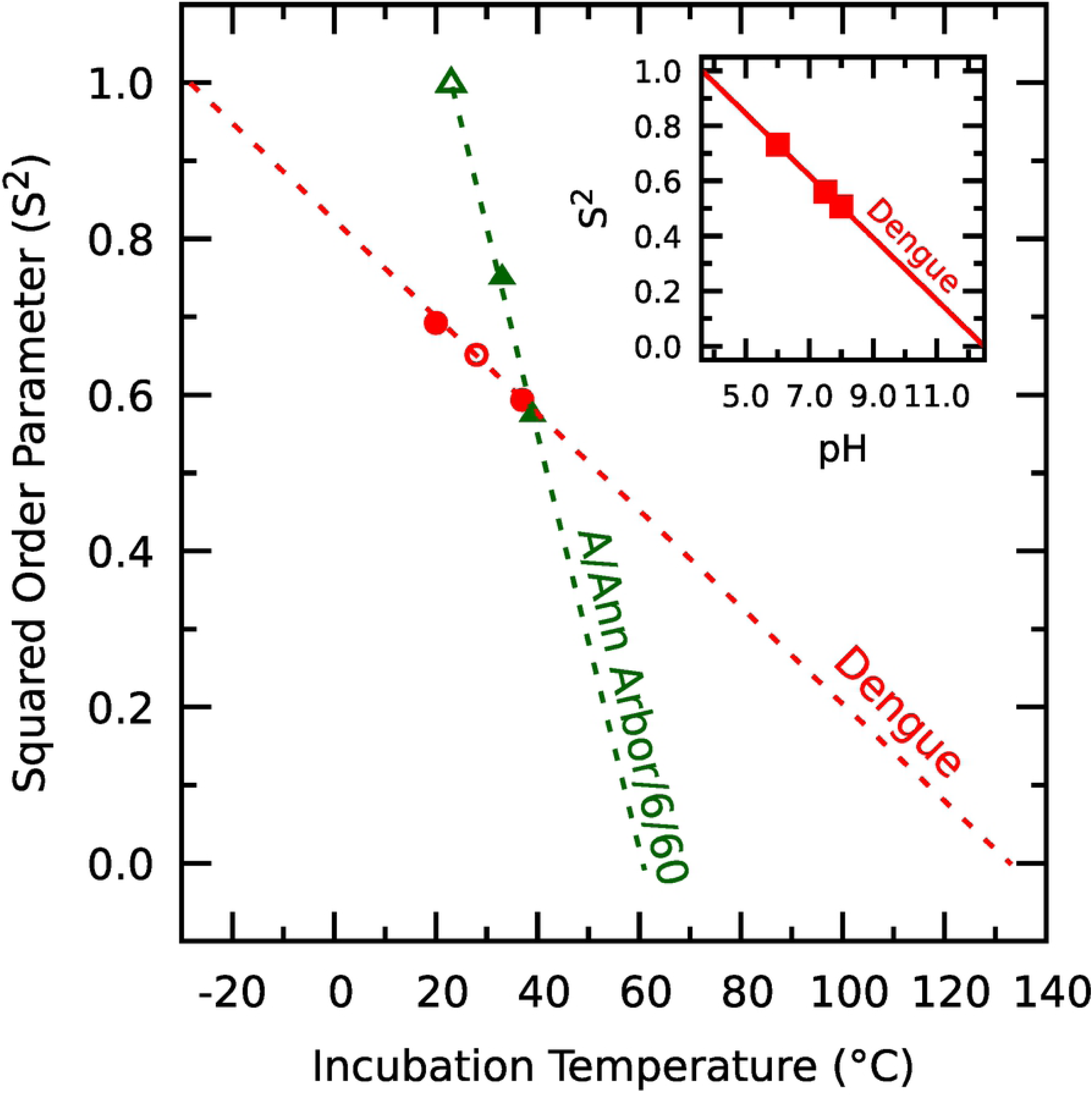
Square of the order parameter as a function of incubation temperature for the Dengue fever virus and the influenza virus. *Inset*: the square of the order parameter as a function of the pH environment of the Dengue fever virus. Data extracted from: filled red circles [26], unfilled red circle [27], filled green triangles [28], unfilled green triangles [29], red squares [30].

Further insight into viruses can be achieved by applying the classic Ising model (*31*) to the viral coat proteins, where the *α* viral coat protein is assigned a spin “up” and the *β* viral coat protein assigned a spin “down.” Such an approach allows us to cast a physical or system property *P* (provided it is dominated by the action of both viral coat proteins) as *P*(*S,x*) *=* [*P*(*S =* 1,0.5) − *P*(*S=0,x*)]*S*^2^ + *P*(*S=*0,*x*). The Ising model equation therefore predicts a linear correlation of the property to the *square* of the order parameter, or *S*^2^. Applying the model to four different virus families (henipaviruses, flaviviruses, influenza viruses and coronaviruses) with the case fatality rate of the virus as the system property, yields remarkable agreement as illustrated in Fig. 5A. It suggests, among other things, that such an approach enables a prediction of the severity of any new virus provided that other members of the same family are well documented. A possible explanation for the linear trends observed in Fig. 5A can be found from within a motif perspective, as we observe that for all four families, as the disorder increases, the fatality rate also increases. Increasing disorder will cause more single-type viral coat protein motifs to form, for example, more hemagglutinin-heavy motifs in the case of influenza, envelope-heavy motifs in flaviviruses, and spike-heavy motifs in coronaviruses. These viral coat proteins are responsible for binding to the hosts cells, so a higher number of such motifs likely translates into a higher probability of the virions successfully attaching and subsequently infecting host cells, and a corresponding higher case fatality rate seen in each family with decreasing *S*^2^ values. It is also interesting to note that while all of the available experimental data points on the Ising model plot in Fig. 5A have a positive value for case fatality rate, extrapolating the lines to the fully ordered (*S* = 1) case yields negative values. Far from being unphysical, we suggest that such values can be explained by considering that the opposite to fatality would be a measure of symbiosis enabled by the virus. An example that supports this interpretation comes from studies of mitochondria (in which the equivalent to viral coat proteins would be the types of porins on its outer surface) and corresponding diseases. As shown in Fig 5B, healthy mitochondria that exist in cells have very high *S*^2^ values, while mitochondria that are responsible for diseases have a lower *S*^2^ value.

**Fig 5.**
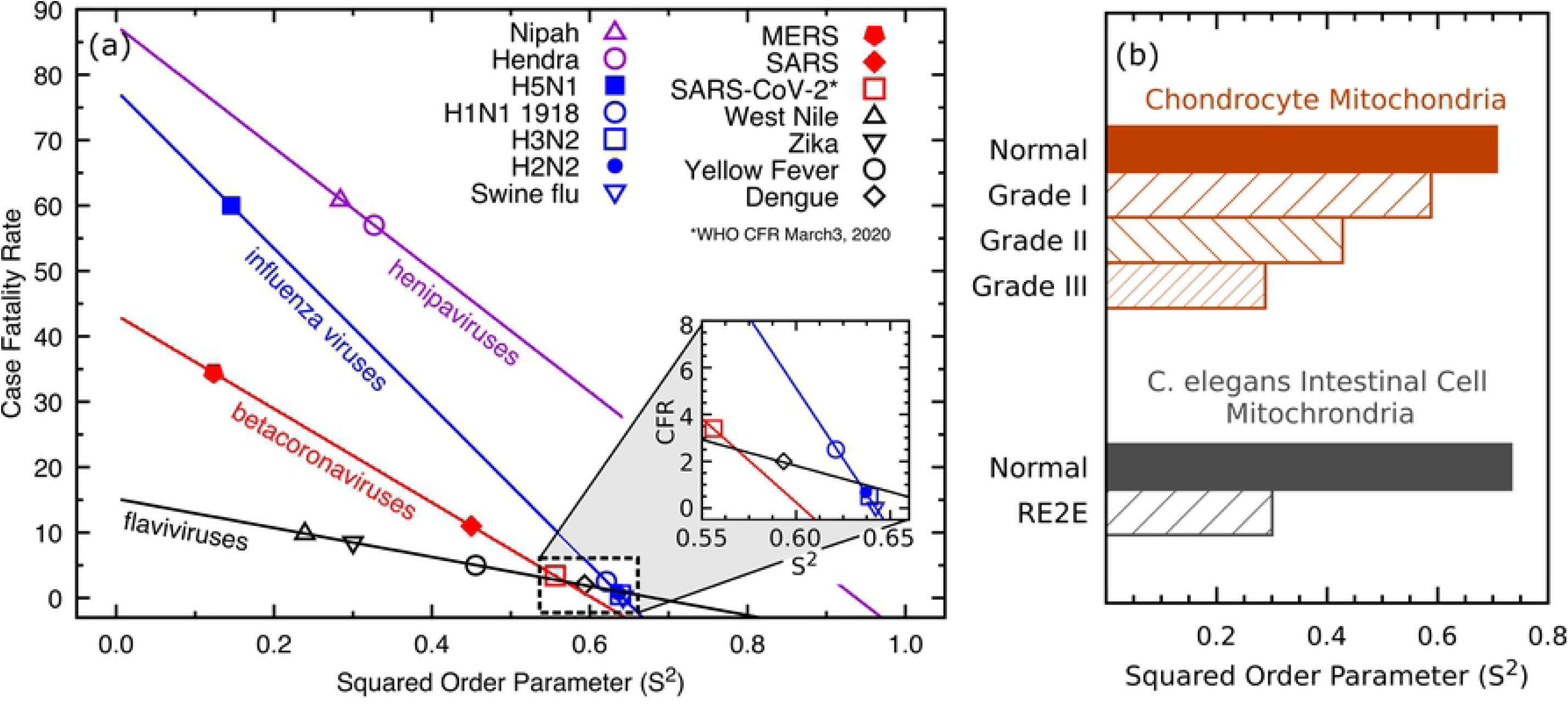
Ising model analysis applied to virus-based diseases. (a) Case fatality rate [1,32-43] as a function of the system order parameter for three viral strains as determined by analysis of published electron microscopy images [15, 26, 44-53]. (b) *S*^2^ values for healthy and disease-causing mitochondria. Data extracted for chondrocyte mitochondria from [54] and for C. elegans mitochondria from [55].

The motif-based disorder perspective also allows for another interesting observation. It is well-known that the 1918 H1N1 virus was deadlier during the second wave in the autumn than during the initial wave of infections during the spring [56]. It is generally held that this second deadlier wave was due to the virus mutating, which can be understood through the lens of motifs and the Ising model. From Fig. 4, we see that increasing the incubation temperature of a virus decreases its order parameter, and from Fig 5, we can see that in general (and specifically for influenza viruses) decreasing *S*^2^ of a given virus increases the mortality rate. Thus, in the case of the 1918 H1N1 pandemic, a possible explanation for the higher autumnal death toll is that the warmer weather created higher incubation temperatures for the virus that caused its order parameter to decrease, leading to an increase in associated mortality rate. Support for such a conclusion can be found in the study by Harding et al., for example, who reported a 0.2 °C increase in the core body temperature from the winter to summer months [57]. Using the linear trends for influenza from Fig. 4, such an incubation temperature change would cause a decrease of 0.0053 in the squared order parameter of the virus, which in turn would correspond to an increase of 0.64 percentage points in the case fatality rate.

## Conclusion

In conclusion, we present a methodology for viewing viruses through the lens of viral coat protein motifs, specifically that disorder and relative fractional composition of viral coat proteins determine the range of viral coat protein structural motifs present on viruses and VLPs used in vaccines. We present a method for quantizing the degree the disorder through an order parameter, *S*, which can be measured using electron microscopy images. Additionally, combining a quantitative measure of disorder with an Ising model allows for deeper insights into the root cause for virus properties in a population, as well as guidance in terms of achieving the characteristics needed for vaccines.

## Materials and methods

The order parameter for lattice structures can be measured using a variety of experimental techniques, such as x-ray diffraction, Raman spectroscopy or electron diffraction [8]. The order parameter of a sample may also be calculated from transmission electron microscopy (TEM) images. In such images, the pixel intensity is less in disordered regions than in ordered regions; this stems from the fact that electrons are incoherently scattered by disordered stacks of atoms as opposed to the coherent diffraction that occurs from well-ordered stacks of atoms [9]. The *S*^2^ value (square of the order parameter) of a sample is equal to the percentage of area corresponding to bright regions. The bright and dark areas corresponding to the ordered and disordered regions can be more easily determined and measured by thresholding the image near the average pixel intensity of the bright regions. The Supplemental Materials section provides evidence for the equivalence between the methods of calculating *S*^2^ from TEM and Raman spectroscopy, specifically surface-enhanced Raman spectroscopy (SERS).

## Acknowledgments

This work stems from studies of disorder in crystalline semiconductor materials, funded in part by the National Science Foundation (DMR-1410915).

## Supporting information

**S1 Fig. Order parameter extraction.** (a) Raman spectroscopy data of an H1N1 VLP extracted from Fig. 4 of ref. [58], along with the fitted ordered and disordered phase peaks. (b) The pixel intensity histogram of the TEM image of the H1N1 VLP extracted from Fig. 3 of ref. [58].

**S1 Table. Order parameter comparison.** Comparison of order parameter (*S*) values extracted from transmission electron microscopy (TEM) and surface-enhanced Raman spectroscopy (SERS) for three different virus-like particles (VLPs), provided as *S*^2^. Data extracted from Lin et al. [58].

